# Divergent methane production and microbial community responses to carbon enrichment in freshwater wetland sediments

**DOI:** 10.64898/2026.11.10.765651

**Authors:** Julia Ghantous, Hassan Azaizah, Elad Levintal, Rotem Golan, Yoav Oved Rosenberg, Gilad Antler, Keren Yanuka-Golub

## Abstract

Methane (CH_4_) accumulation in wetland sediments reflects interactions among organic matter, sediment chemistry, and microbial communities. We combined depth-resolved field characterization across four wetland habitats in northern Israel with complementary incubations to examine methane accumulation and responses to labile-carbon enrichment. Sediment cores were characterized using Rock-Eval analysis, porewater chemistry, oxygen profiling at three sites, and *mcrA* amplicon sequencing. High CH_4_ concentrations occurred under contrasting conditions: Ein Nimfit (EN) contained approximately 12% total organic carbon and reached 2.1 mM CH₄, whereas Ein Afek site EA2 contained approximately 5% carbon and accumulated 0.7–1.7 mM CH_4_. Both showed sulfate depletion, but EN exhibited pronounced sulfide accumulation, whereas EA2 showed greater dissolved iron and nutrient enrichment. Their classified methanogenic assemblages also differed. EA1 remained methane poor despite increasing carbon content with depth, while EA3 combined low carbon content and CH_4_ accumulation with detectable methanogens and lower methanogenic diversity. In a 36-day bottle experiments using sediments from two of the four wetland habitats, sucrose-amended EA3 exhibited a transient maximum of 18.8 mmol collected CH_4_ kg⁻¹ wet sediment during Days 22–29, whereas EN reached its highest collected amount during Days 29–36. Methane proportions increased before the large output increase in EA3 but remained comparatively stable and high in EN. Soluble chemical oxygen demand trajectories and microbial community changes also differed. These findings identify distinct biogeochemical settings supporting methane accumulation and demonstrate methanogenic potential in low-CH_4_ sediment, emphasizing the complementary information provided by field profiles, microbial characterization, and exploratory enrichment experiments.

**Importance:** Wetlands store carbon but also produce methane, making it important to understand why methane accumulates in some sediments and remains scarce in others. Across four habitats in northern Israel, methane-rich sediments differed in their carbon content, dissolved chemical constituents, and methane-producing archaeal communities. Conversely, carbon-rich layers and the presence of methane producers did not consistently coincide with methane accumulation. Complementary laboratory incubations showed that sediment containing little methane in the field could express substantial methane-production potential after labile carbon amendment, accompanied by changes in bacterial and archaeal communities. The study brings together sediment properties, chemical profiles, and microbial responses to distinguish the conditions associated with methane accumulation from the capacity to produce methane. This distinction helps frame future investigations of how changing carbon inputs and wetland conditions influence microbial carbon processing.

## 1. Introduction

Wetlands are among the most biologically productive ecosystems on Earth and play an important role in global carbon cycling by storing large amounts of organic carbon under water-saturated conditions. Limited oxygen (O_2_) diffusion in wetland soils promotes anaerobic biogeochemical processes that slow organic matter (OM) decomposition and facilitate long-term carbon accumulation (Kayranli et al., 2010; Reddy et al., 2022). As a result, wetlands store an estimated ∼600 Pg of carbon as soil organic matter (SOM), representing a substantial fraction of the global terrestrial carbon stocks (Goldstein et al., 2020). However, these ecosystems are also major natural sources of methane (CH_4_), a potent greenhouse gas (GHG) produced during anaerobic OM mineralization.

Wetland degradation and land-use changes can destabilize stored carbon pools, leading to increased GHG emissions (Biancalani et al., 2014; Deshmukh et al., 2021; Frolking et al., 2011; Tapio-Biström et al., 2012). Although wetland restoration is widely promoted as a climate mitigation strategy, restored systems may function as net sources of CH_4_, particularly during the early stages of recovery or under poorly constrained hydrological and biogeochemical conditions (Lorenz & Lal, 2018). Understanding wetland CH₄ dynamics therefore requires examining the processes that govern sediment carbon turnover and methane production under anaerobic conditions.

Carbon cycling in wetlands is primarily mediated by microorganisms that decompose OM and drive greenhouse gas production through tightly coupled metabolic networks (Bechtold et al., 2025). Under O_2_ limitation, microbial OM oxidation is traditionally described as proceeding through a thermodynamic sequence of anaerobic respiratory pathways in which microorganisms utilize alternative electron acceptors, such as nitrate, Mn(IV), Fe(III), and sulfate, before terminal carbon mineralization is coupled to methanogenesis (Alewell et al., 2008; Laanbroek, 2010; Megonigal et al., 2003; Pester et al., 2012; Reddy et al., 2022). In this redox framework, CH_4_ is produced through multiple methanogenic pathways, including hydrogenotrophic methanogenesis by carbon dioxide (CO_2_) reduction, acetoclastic methanogenesis via acetate cleavage, and methylotrophic or hydrogen-dependent methylotrophic methanogenesis from methylated compounds, such as methanol, methylamines, and methylated sulfur compounds (Bueno De Mesquita et al., 2023; Hartman et al., 2024). These pathways represent the terminal stages of anaerobic carbon mineralization and occur as free-energy yield declines across terminal electron-accepting processes (Conrad, 2020; Ferry, 2010, 2011). However, natural wetland sediments rarely follow this sequence as a simple linear progression. Instead, redox processes may overlap spatially and temporally owing to microscale heterogeneity, fluctuating hydrology, variable OM quality, diffusion of electron acceptors, plant-mediated O_2_ transport, and microbial syntrophic interactions (Dalcin Martins et al., 2017; Liu et al., 2025; Reiche et al., 2008; Sela-Adler et al., 2017).

The balance between these competing metabolic pathways plays a central role in regulating CH_4_ production. In addition to redox constraints, the availability, composition, and reactivity of OM substrates strongly influence microbial metabolism and carbon mineralization. Biological OM lability refers to the susceptibility of organic carbon to microbial degradation, supporting microbial respiration and methanogenesis under prevailing environmental conditions (Scott et al., 2020; Yarwood, 2018; Zheng et al., 2025). The bioavailability of OM to microorganisms depends on several factors, including its molecular composition and complexity, solubility, accessibility to extracellular enzymes, association with mineral surfaces, and environmental conditions that regulate microbial degradation (Lehmann & Kleber, 2015; Marschner & Kalbitz, 2003; Schmidt et al., 2011).

Geochemical characterization complements assessments of biological OM lability by providing operational measures of sediment OM composition and thermal stability. Rock–Eval analysis characterizes OM through programmed heating, during which less thermally stable fractions decompose at lower temperatures. Thermal lability may be associated with susceptibility to microbial degradation, although this relationship depends on OM composition and environmental conditions (Disnar et al., 2003; Gerera et al., 2025; Sebag et al., 2016). Rock–Eval measurements therefore describe thermal decomposition rather than directly measuring microbial bioavailability (Hazera et al., 2026; Sapir et al., 2026). These two concepts may be related but are not necessarily equivalent, because thermally labile OM may not be equally accessible or metabolizable by anaerobic microbial communities *in-situ*. Biologically labile carbon inputs, including plant-derived root exudates, can stimulate microbial activity and enhance methanogenesis either directly or through priming effects (Waldo et al., 2019).

Previous studies in freshwater lake and reservoir sediments have linked variation in CH_4_-production rates to organic matter supply and composition, as well as microbial community composition (Berberich et al., 2020; Bertolet et al., 2019). However, these relationships do not establish how field CH_4_ conditions relate to sediment responses to fresh organic-carbon inputs. A key question is how sediments with contrasting field CH_4_ concentrations and methanogen communities respond to the same labile-carbon addition, particularly in the magnitude and timing of methane production and the accompanying changes in the broader microbial community. To address this question, we investigated four wetland habitats in northern Israel: three ponds within the Ein Afek Nature Reserve wetland complex (EA1–EA3) and Ein Nimfit wetland (EN). We combined depth-resolved measurements of sediment OM properties and porewater chemistry, including CH_4_ concentrations, with characterization of methanogen communities using *mcrA* amplicon sequencing. Controlled anaerobic incubations of sediments from EA3 and EN examined methane-production responses to labile-carbon addition and accompanying changes in bacterial and archaeal communities using 16S rRNA gene amplicon sequencing. We hypothesized that field CH_4_ concentrations alone would not predict the magnitude or timing of methane-production responses to labile-carbon addition, and that contrasting responses would be accompanied by distinct changes in microbial community composition.

## 2. Methods

### 2.1 Study sites

We investigated sediment biogeochemistry and CH_4_ dynamics in two spring-fed wetlands within the Na‘aman River basin in northern Israel: Ein Afek Nature Reserve and Ein Nimfit (Fig. 1A). Three habitats within Ein Afek (EA1–EA3) and one location within Ein Nimfit (EN) were studied. Both wetlands have experienced hydrological changes associated with groundwater pumping and water management. Ein Afek comprises a network of spring-fed ponds with a main outlet to the Na‘aman River. Groundwater pumping contributed to the cessation of spring flow during 2008–2012, when supplementary water was supplied from a borehole; substantial natural spring flow resumed in 2013. Ein Nimfit comprises a natural spring-fed pond and an artificial pond excavated in 2000, and its water regime has been affected by diversion of spring water to adjacent fishponds. Historical chloride concentrations were reported as 600–1,100 mg Cl⁻ L⁻¹ at Ein Afek and 800–1,000 mg Cl⁻ L⁻¹ at Ein Nimfit, the latter based on monitoring during 2010–2012, consistent with historically brackish source water at both wetlands (Sapir and Elron, 2014).

**Figure 1.**
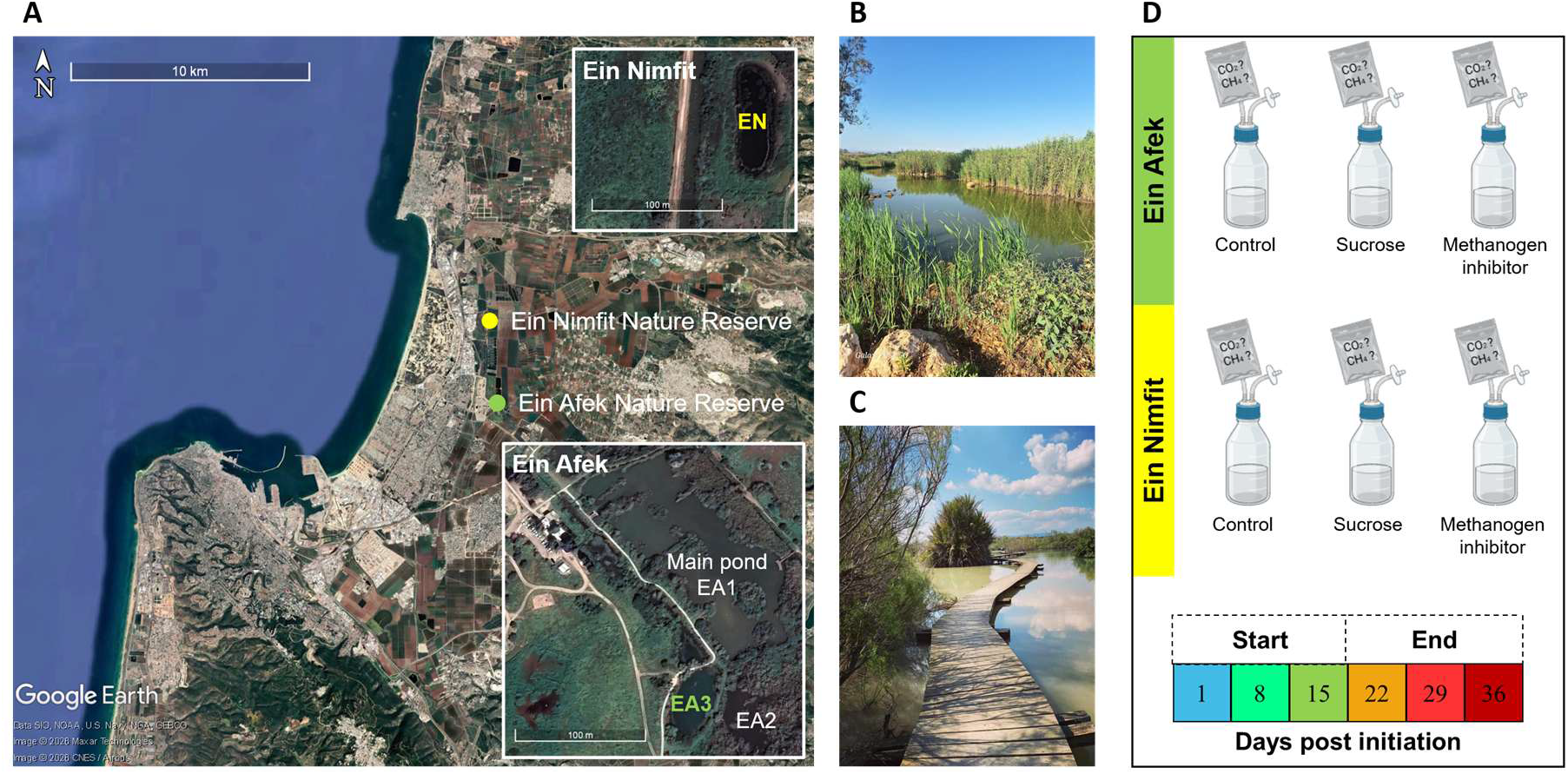
Study sites: (A) Map showing the locations of Ein Afek Nature Reserve (EA) and Ein Nimfit Nature Reserve (EN) in northern Israel, with enlarged views of the sampling sites at Ein Nimfit (EN) and Ein Afek (EA1–EA3). (B) Photograph of the Ein Nimfit wetland. (C) Photograph of the Ein Afek wetland. Satellite imagery was obtained from Google Earth. (D) Bottle incubation experimental design.

Depth-resolved sediment cores were collected to characterize sediment organic matter properties, porewater chemistry, CH_4_ concentrations, and methanogen community composition using *mcrA* amplicon sequencing. A separate anaerobic bottle-incubation experiment using sediments from EA3 and EN examined methane-production responses to labile-carbon addition and accompanying changes in bacterial and archaeal community composition using 16S rRNA gene amplicon sequencing.

### 2.2 Sediment core collection and analyses

Intact vertical sediment cores were collected in September 2024 from four locations: EA1, EA2, and EA3 within the Ein Afek Nature Reserve complex and EN within Ein Nimfit (Fig. 1A). The recovered core depths differed among sites, reaching 35 cm at EA1, 28.5 cm at EA2, 12 cm at EA3, and 27.5 cm at EN. Porewater and sediment were sampled at approximately 3 cm intervals throughout the recovered depth of each core. Porewater was collected using Rhizon samplers inserted into the cores, while sediment subsamples were obtained through pre-made sampling ports in the core liners using a 3 mL syringe. Sediment subsamples were divided into separate aliquots for organic carbon analysis, dissolved CH_4_ quantification, and DNA extraction followed by *mcrA* amplicon sequencing.

### 2.3 Biogeochemical analyses of core samples

#### 2.3.1 Sediment organic carbon quantity and thermal indices

Sediment organic carbon (SOC) was characterized by Rock-Eval 7S thermal analysis (RE-7S, Vinci Technologies) using a two-stage pyrolysis and oxidation procedure. Approximately 20 mg of dried, homogenized sediment was analyzed for total organic carbon (TOC), mineral carbon (MINC), and thermal indices describing OM composition. Rock-Eval parameters included the hydrogen index (HI), oxygen index (OI), and Tmax. In addition, I and R indices were calculated from the S2 thermogram to represent the relatively thermally labile and refractory fractions of SOC, respectively, as described by Sapir et al., 2026 and Sebag et al., 2016.

#### 2.3.2 Porewater solute analysis

Porewater was extracted from the sediment cores at approximately 3 cm intervals using Rhizon samplers. Ammonium (NH_4_^+^) and phosphate (PO_4_^3-^) were measured using a continuous-flow nutrient analyzer (Gallery Discrete Analyzer, Thermo Fisher Scientific). Dissolved ferrous iron (Fe^2+^) was determined using the ferrozine method (Stookey, 1970), and sulfide (H_2_S) was measured using a modified methylene blue assay (Cline, 1969). Major cations were analyzed using microwave plasma atomic emission spectrometry (MP-AES 4210, Agilent Technologies).

#### 2.3.3 Dissolved inorganic carbon and carbon isotopes

Porewater dissolved inorganic carbon (DIC) concentrations and the stable carbon isotopic composition of dissolved inorganic carbon (δ^13^C-DIC) were measured to characterize carbon mineralization and depth-dependent isotopic variation. Total alkalinity was determined by acid–base titration with 0.005 M HCl using an automated titrator (TitroLine 7000, SI Analytics) and calculated using a Gran plot. DIC concentrations were measured using an AS-C6L DIC analyzer (Apollo SciTech), and δ^13^C-DIC was measured using a GasBench II extraction system coupled to a DELTA V isotope-ratio mass spectrometer (Thermo Fisher Scientific).

#### 2.3.4 Dissolved CH_4_ concentration in sediment porewater

Dissolved CH_4_ concentrations were determined from NaOH-preserved sediment subsamples using headspace analysis (Liu et al., 2025). Headspace CH_4_ was quantified using a gas analyzer (7810, LI-COR) equipped with a small-volume sampling kit. Headspace CH_4_ concentrations were converted to dissolved concentrations using the ideal gas law and normalized to the porewater volume of each sediment subsample. Final CH_4_ concentrations were expressed as mM. Detailed sample preservation, headspace sampling, replicate measurements, and concentration calculations are provided in the Supplementary Information.

#### 2.3.5 Dissolved O_2_ profiles in sediment porewater

Dissolved O₂ profiles were measured in smaller intact sediment cores collected specifically for O₂ profiling, separately from the cores used for sediment and porewater analyses, using an optical O_2_ microsensor (PyroScience GmbH) with a tip diameter of 230 µm. The sensor was calibrated using O_2_-saturated water (100% air saturation), and the overlying water was removed shortly before measurement. Vertical profiles were measured at 0.2 mm intervals from the sediment surface to approximately 2 cm depth to capture the steep O_2_ gradients near the sediment–water interface, which required a finer spatial resolution than the ∼3 cm intervals used for sediment and porewater sampling. Measurements were performed using a micromanipulator, and O_2_ concentrations were expressed as percentages of air saturation. Profiles were obtained for EA1, EA2, and EN; O_2_ profiling was not performed at EA3 because an intact core suitable for microsensor measurements could not be recovered.

### 2.4 Bottle incubation experiment

Bottle incubations were conducted using sediments from EA3 and EN to examine gas-production and microbial community responses to labile-carbon addition and methanogenesis inhibition. Each bottle contained 1,200 g wet sediment and 1,000 g medium. Three treatments were established for each sediment: unamended control, sucrose addition, and sucrose + sodium 2-bromoethanesulfonate (BES; 40 mM). Each sediment–treatment combination was represented by one bottle, yielding six unreplicated experimental units. Bottle headspaces were flushed with N₂, and incubations were maintained at 30°C for 36 days. Sucrose was supplied initially at 0.1% w/w of the sediment–medium mixture, equivalent to 2.2 g per bottle, and replenished when measured soluble chemical oxygen demand (sCOD) fell below 300 mg O_2_ L^-1^. Feeding schedules consequently differed between bottles. For sCOD analysis, liquid samples were filtered through 0.45-µm membranes, digested at 150°C for 2 h, and measured spectrophotometrically at 620 nm using a calibration curve prepared from standards of known COD.

Gas measurements and sediment sampling for microbial community analysis were performed on Days 8, 15, 22, 29, and 36. Full gas-bag volumes and measured CH_4_ and CO_2_ fractions were used to calculate collected gas amounts using the ideal gas law. Amounts represented successive seven-day collection intervals and were normalized to initial wet sediment mass. Gas bags were emptied after each measurement. Methane was additionally expressed as a percentage of combined measured CH_4_ and CO_2_ to characterize gas composition while reducing the influence of common dilution by other gases during sampling. Three technical DNA-extraction replicates were processed per sediment sample. For descriptive community comparisons, sampling points were grouped retrospectively into early (Days 8–15) and late (Days 22–36) phases based on the observed gas-collection patterns. Gas and community trajectories were interpreted as exploratory observations because bottles were unreplicated. Detailed incubation procedures, feeding records, analytical methods, gas calculations, and documented sealing and sampling issues are provided in the Supplementary Information. Medium composition is provided inYusim et al., (2025) and Table S1.

### 2.5 DNA extraction, PCR amplification, and amplicon sequencing

DNA was extracted from environmental core and bottle-incubation samples using the DNeasy PowerSoil Pro Kit (QIAGEN), with the two sample sets processed separately. Environmental core samples were analyzed by *mcrA* amplicon sequencing to characterize methanogenic communities (Luton et al., 2002), whereas bottle-incubation samples were analyzed by 16S rRNA gene (V3–V4) amplicon sequencing to characterize bacterial and archaeal communities (Naqib et al., 2018; Takahashi et al., 2014). Libraries were sequenced on an Illumina MiSeq platform at the DNA Services Facility, University of Illinois Chicago, USA. Detailed PCR mixtures, primer sequences, and thermal cycling conditions are provided in Tables S2–S4. Raw *mcrA* and 16S rRNA gene sequences were deposited in the NCBI Sequence Read Archive under accession numbers PRJNA1537901 and PRJNA1354830, respectively.

### 2.6 Sequence data processing

Raw reads were quality-checked using FastQC, merged using PEAR (Zhang et al., 2014), and processed in QIIME 2 (Bolyen et al., 2019), with amplicon sequence variants (ASVs) inferred using DADA2 (Callahan et al., 2016). The *mcrA* and 16S rRNA gene datasets were processed separately, and the resulting tables were used for downstream community analyses in R. Detailed processing parameters and filtering criteria are provided in the Supplementary Information.

#### 2.6.1 Methanogens sequencing from sediment cores

A total of 34 sediment subsamples from the four cores were analyzed for *mcrA* community composition (EN, n = 7; EA1, n = 10; EA2, n = 11; EA3, n = 6), with sample numbers reflecting differences in recoverable core length. Processing was performed using QIIME 2 version 2023.5. Primer sequences were removed using Cutadapt (Martin, 2011) before DADA2 denoising. Following quality filtering, 32 samples were retained for downstream analysis. The ASV table was filtered and normalized using cumulative sum scaling (CSS). Community composition was evaluated using Bray-Curtis dissimilarities, nonmetric multidimensional scaling (NMDS), and PERMANOVA. Associations with sediment geochemical properties were explored using environmental vector fitting. Primer sequences, detailed normalization and statistical procedures are provided in the Supplementary Information.

#### 2.6.2 Prokaryotic sequencing and community analysis of the bottle experiment

ASVs were taxonomically assigned against the SILVA database. Alpha diversity was evaluated using Shannon and Simpson diversity indices and observed ASV richness. Community dissimilarities were assessed using Bray-Curtis distances and visualized using NMDS. Taxonomic composition was examined at the family and genus levels, and selected taxa were assigned to putative functional guilds based on taxonomy. Associations between microbial taxa, putative functional guilds, and CH_4_ and CO_2_ production were explored using Spearman correlations. Because each site–treatment combination was represented by one bottle, community patterns and correlations were interpreted as exploratory. Primer sequences, detailed normalization, diversity analyses, functional guild assignments, and correlation procedures are provided in the Supplementary Information.

## 3. Results

### 3.1 Sediment organic carbon quantity and composition differed among wetland sites

To characterize differences in sediment carbon properties among the studied sites, we first examined depth profiles of TOC, MINC, and the Rock-Eval-derived HI and OI indices (Fig. 2). TOC differed markedly among sites (Fig. 2A), with EN showing the highest organic carbon content (∼12%) across most of the sediment profile, EA2 showing intermediate values (∼5%), and EA3 remaining consistently low (∼1%). EA1 showed a distinct vertical pattern, with TOC increasing with depth and reaching approximately 8% in the deeper sediment.

**Figure 2.**
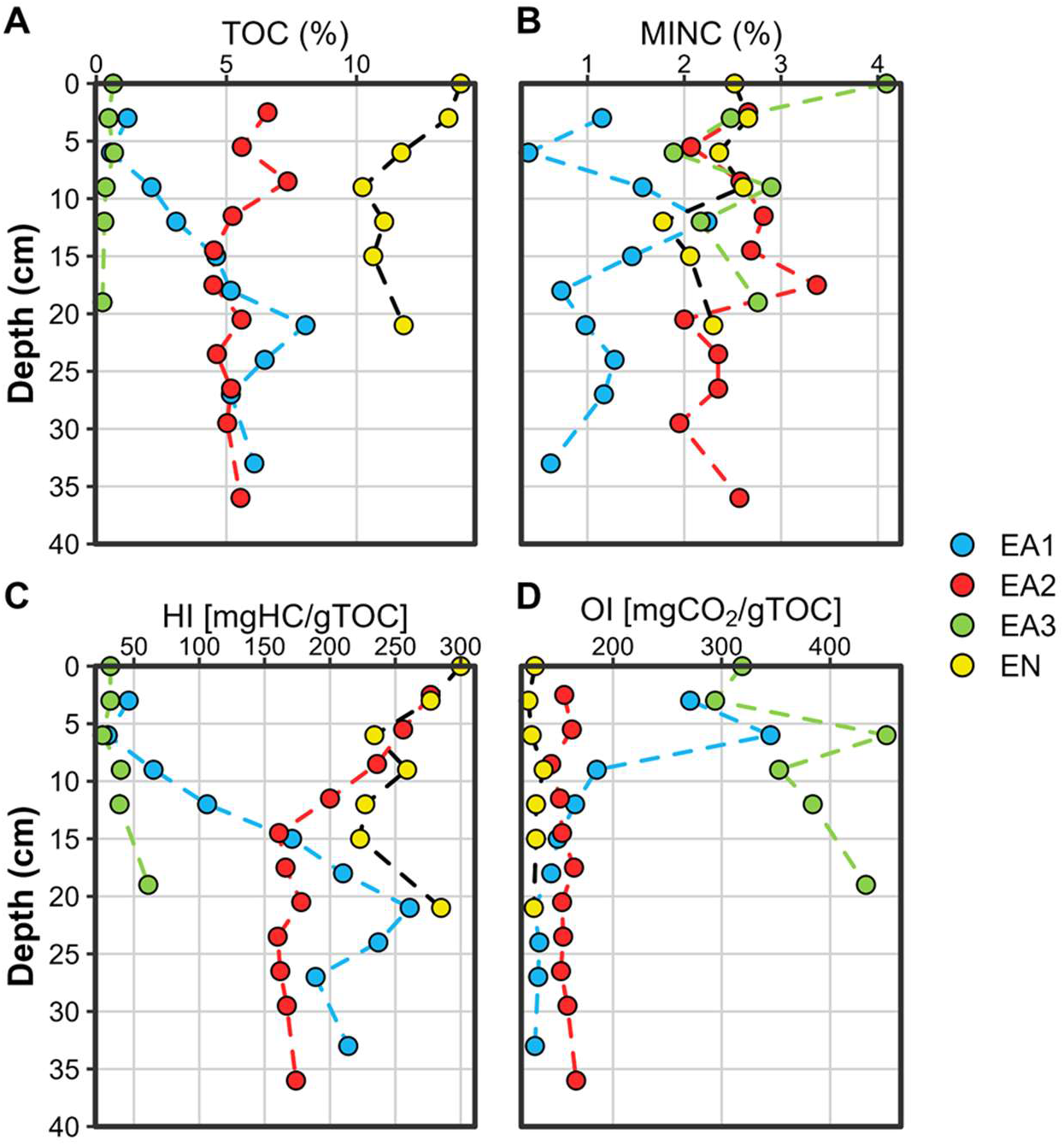
Depth profiles of sediment carbon content and Rock-Eval-derived organic matter indices. (A) Total organic carbon (TOC, %); (B) mineral carbon (MINC, %); (C) hydrogen index (HI, mg HC g⁻¹ TOC); and (D) oxygen index (OI, mg CO₂ g⁻¹ TOC) in sediments from Ein Afek Nature Reserve sites EA1-EA3 and Ein Nimfit (EN). HI and OI characterize the hydrogen and oxygen contents of sediment organic matter, respectively.

The sites also differed markedly in their Rock-Eval-derived OM characteristics (Fig. 2B-D). In the upper sediment, EN and EA2 showed relatively similar HI and OI values. EN maintained relatively high HI and low OI throughout the profile, whereas EA2 showed a gradual decrease in HI with depth. EA1 exhibited a pronounced depth-dependent shift, with increasing TOC and HI accompanied by decreasing OI. In contrast, EA3 was characterized by consistently low TOC and HI, and substantially higher OI values throughout the profile.

Sediment OM differed markedly among the sampled cores. EN and EA3 showed the strongest contrast: EN had high TOC & HI and low OI, whereas EA3 showed the opposite pattern. EA1 exhibited a pronounced depth gradient, with TOC and HI increasing and OI decreasing in deeper sediments.

Porewater chemistry and dissolved CH_4_ concentrations varied markedly among cores and with depth (Fig. 3). CH_4_ concentrations were highest in EN, reaching approximately 2.1 mM, followed by EA2, where values ranged from approximately 0.7 to 1.7 mM. In both cores, CH_4_ increased below approximately 5 cm, coinciding with pronounced porewater SO_4_²⁻ depletion. EA1 concentrations remained near zero, whereas EA3 showed generally low concentrations with a localized subsurface maximum of approximately 0.5 mM.

**Figure 3.**
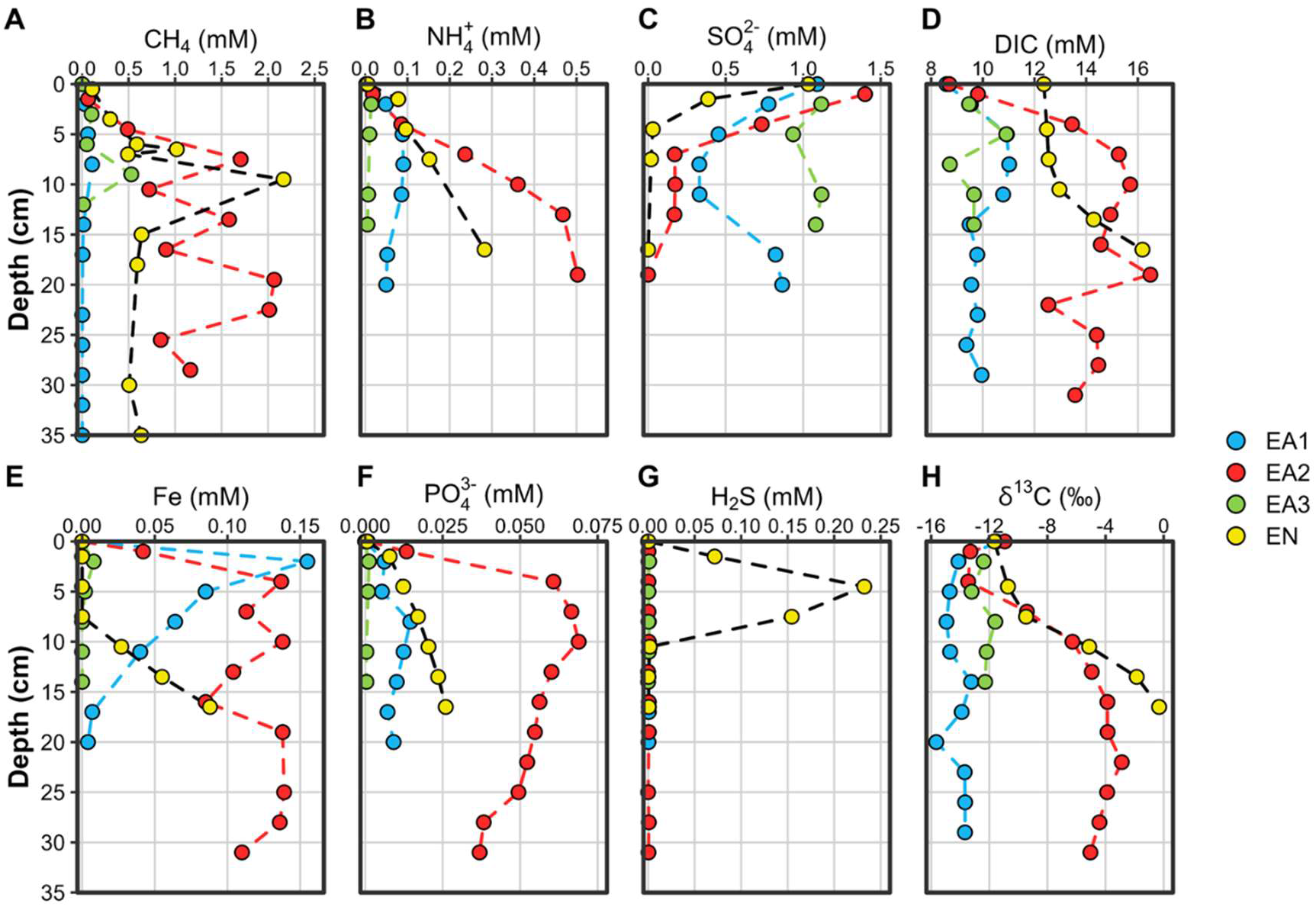
Depth profiles of dissolved CH₄ and porewater biogeochemistry. (A) CH_4_; (B) NH_4_⁺; (C) SO_4_²⁻; (D) dissolved inorganic carbon (DIC); (E) dissolved Fe; (F) PO_4_³⁻; (G) H_2_S; and (H) stable carbon isotopic composition of DIC (δ¹³C-DIC) in sediment porewater from Ein Afek sites EA1-EA3 and Ein Nimfit (EN). Concentrations are expressed in mM and δ¹³C-DIC in ‰. CH_4_ values represent dissolved porewater concentrations rather than atmospheric emission fluxes.

The CH_4_-enriched EA2 and EN sites were also characterized by distinct porewater biogeochemical patterns. EA2 showed pronounced increases in NH_4_⁺ and DIC with depth (Fig. 3A), with NH_4_⁺ reaching approximately 0.5 mM and DIC approximately 15–16 mM in the deeper sediment (Fig. 3B). PO_4_^3-^ was also substantially higher in EA2 than at the other sites, reaching approximately 0.07 mM in the shallow sediment and remaining elevated with depth (Fig. 3F). In contrast, EN was distinguished by a pronounced subsurface H_2_S maximum, reaching approximately 0.23 mM at ∼5 cm and declining sharply below this depth (Fig. 3G). Dissolved Fe remained low within this H_2_S-enriched interval and increased at greater depth, whereas EA2 maintained comparatively high Fe concentrations over much of its profile (Fig. 3E). Thus, although both EN and EA2 showed substantial CH_4_ accumulation following SO_4_^2-^ depletion, their associated porewater profiles were distinct.

DIC profiles showed distinct depth-dependent patterns among sites (Fig. 3D). EA2 displayed a pronounced subsurface increase in DIC between approximately 5 and 20 cm, reaching ∼16 mM, whereas EN reached also ∼16 mM but showed a more gradual increase with depth. Similarly, EA1 and EA3 reached comparable maximum concentrations (∼11 mM), although EA1 showed a clearer increase with depth, while EA3 exhibited relatively limited vertical variation. δ^13^C-DIC values were relatively negative across all four sites (Fig. 3H). With increasing depth, EA2 and EN shifted toward more positive values, reaching approximately −4‰ and 0‰, respectively. In contrast, EA1 and EA3 remained comparatively ^13^C-depleted throughout most of their profiles, with values remaining around −16‰ and −12‰, respectively.

### 3.2 Sediment oxygen profiles and multivariate biogeochemical patterns

Dissolved O₂ profiles showed differences in the depth of the oxic sediment layer among sites. O₂ saturation declined rapidly with depth, reaching near-zero values at approximately 0.5 cm in EN, 0.8 cm in EA2, and 1.7 cm in EA1 (Fig. 4A).

**Figure 4.**
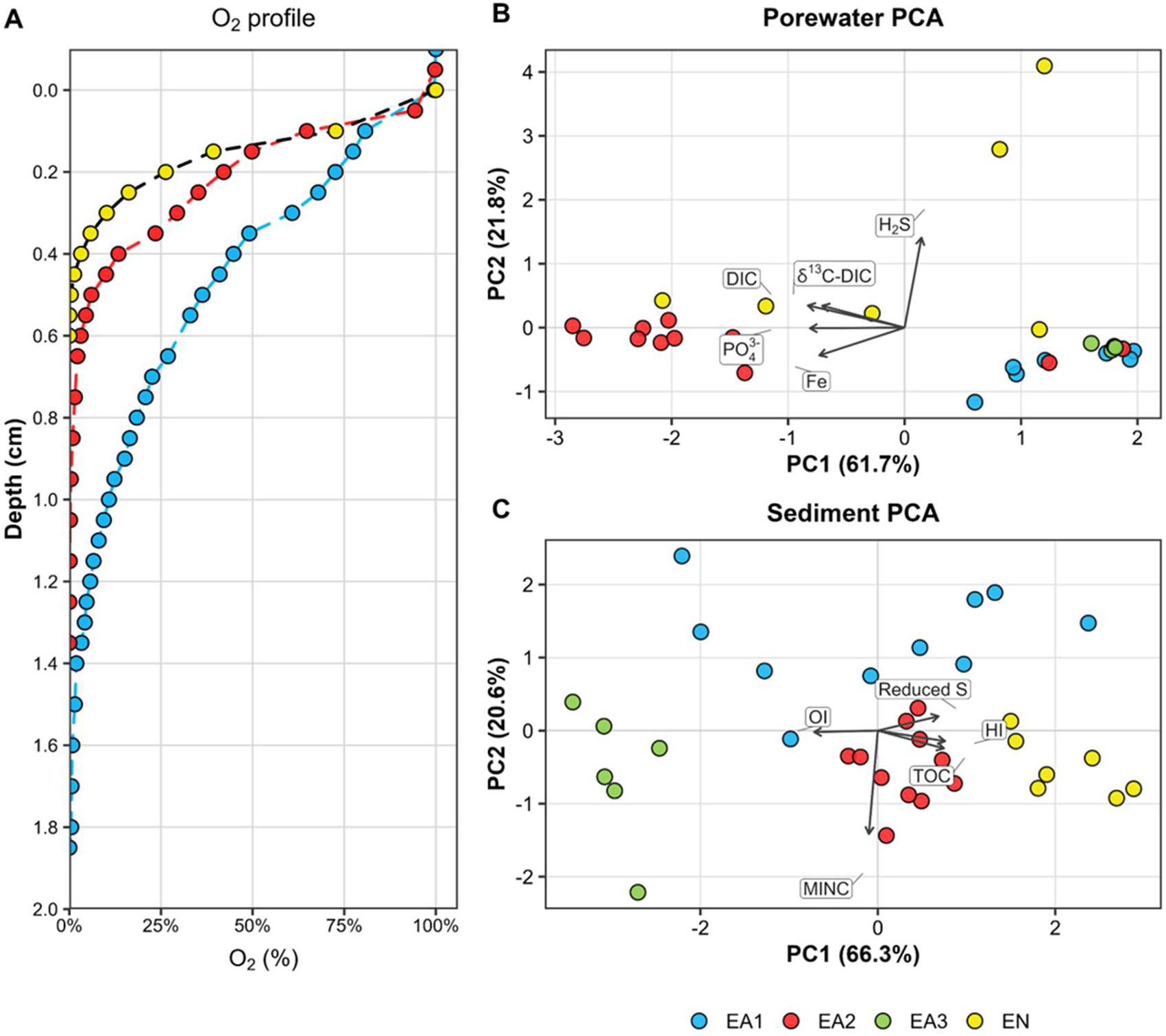
Oxygen distribution and multivariate biogeochemical structure across wetland sediments. (A) Dissolved oxygen (DO, % saturation) microprofiles measured in intact short sediment cores from EA1, EA2, and EN. (B) Principal component analysis (PCA) of porewater chemistry based on H_2_S, dissolved Fe, PO_4_^3-^, DIC, and δ^13^C-DIC. (C) PCA of sediment properties based on TOC, mineral carbon (MINC), hydrogen index (HI), oxygen index (OI), and reduced sulfur. Variables were centered and scaled prior to PCA. Points in the PCA plots represent depth-resolved samples and are colored by site.

PCA of porewater chemistry distinguished the sampled cores (Fig. 4B). PC1 explained 61.7% of the variation and was primarily associated with DIC, PO_4_^3-^, Fe, and δ¹³C-DIC. EA2 samples were positioned predominantly on the negative side of PC1, toward the DIC, PO_4_^3-^, and Fe loadings, whereas EA1 and EA3 generally occupied the positive side. PC2 explained an additional 21.8% of the variation and was strongly associated with H_2_S. EN samples showed the greatest spread along PC2, with some extending toward the H₂S loading, consistent with the subsurface sulfide maximum in the porewater profile (Fig. 3).

PCA of sediment properties captured the contrasting OM characteristics among cores (Fig. 4C). PC1 explained 66.3% of the variation, with TOC, HI, and reduced sulfur loading in the opposite direction to OI. EN and EA3 occupied contrasting positions along this axis: EN samples were associated with TOC, HI, and reduced sulfur, whereas EA3 samples were associated with OI. EA2 occupied an intermediate position. PC2 explained a further 20.6% of the variation and was primarily associated with MINC, distinguishing EA1 mainly along this axis. Together, the sediment and porewater analyses identified contrasting biogeochemical conditions in EN and EA3, complementing their different dissolved CH_4_ profiles (Fig. 3).

### 3.3 Methanogenic community structure across depth profiles

The methanogenic community composition varied among cores and with sediment depth, revealing site-specific structuring of *mcrA* assemblages (Fig. 5). Shannon diversity also differed among cores and was lowest in EA3 among the sampled cores (Fig. S1). Unresolved genus-level assignments dominated the *mcrA* dataset, accounting for an average of 70.0% of relative abundance across samples (range, 39.4–91.4%). Genera displayed individually accounted for 24.8% on average, while low-abundance genera grouped as “Other (<1%)” accounted for 5.2%. The contribution of individually displayed genera varied among cores, averaging 39.7% in EA2, 19.1% in EA1, 17.0% in EN, and 12.0% in EA3.

**Figure 5.**
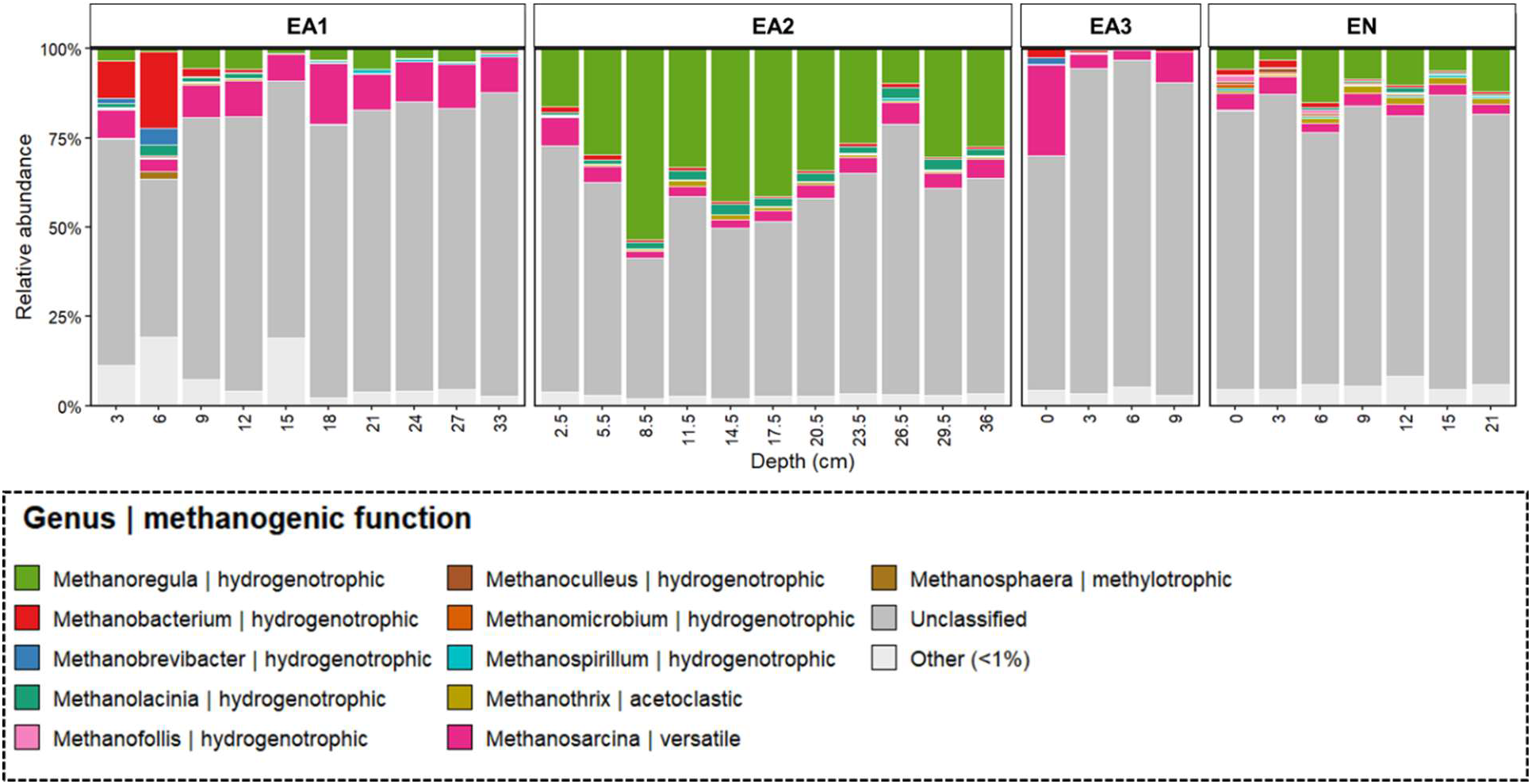
Depth-resolved taxonomic composition of *mcrA* assemblages across wetland sediment cores. Bars show relative abundances calculated from CSS-normalized *mcrA* counts at individual sediment depths in Ein Afek cores EA1–EA3 and Ein Nimfit (EN). Genera reaching ≥1% relative abundance in at least one sample are shown individually. Unclassified, uncultured, and other unre-solved genus-level assignments are grouped as “Unclassified”; remaining low-abundance genera are grouped as “Other (<1%).” CSS, cumulative sum scaling.

EA2 was characterized by a high relative abundance of *Methanoregula*, which averaged 31.4% of the *mcrA* dataset and reached 53.4% at 8.5 cm. This genus remained abundant across much of the profile but declined in the deepest sediments. *Methanosarcina* and *Methanolacinia* occurred at lower relative abundances, averaging 4.2% and 2.2%, respectively. EA1 showed pronounced vertical variation among classified genera. *Methanobacterium* reached 10.5% at 3 cm and 21.5% at 6 cm before declining sharply with depth. In contrast, *Methanosarcina* occurred throughout much of the profile and became more abundant in deeper sediments, reaching 17.2% at 18 cm and approximately 10–12% at several deeper sampling depths. *Methanoregula* occurred at several depths but remained less abundant than in EA2. In EN, *Methanoregula* was the most abundant classified genus, averaging 8.7% and reaching 15.1% at 6 cm. *Methanosarcina* and *Methanothrix* averaged 3.6% and 1.4%, respectively, alongside smaller contributions from other methanogenic genera. EA3 had the highest proportion of unresolved assignments, averaging 83.9%. Among its classified genera, *Methanosarcina* reached 25.4% in the surface sediment but declined substantially below the surface. *Methanoregula* was nearly absent, averaging 0.2%, and the remaining classified genera occurred at low relative abundances.

Overall, EA2 had a strong contribution from *Methanoregula*, EA1 showed pronounced vertical shifts among classified genera, EN contained several classified methanogenic genera, and EA3 had lower Shannon diversity and predominantly unresolved genus-level assignments. The contrasting *mcrA* assemblages, sediment properties, and porewater chemistry of EA3 and EN provided a basis for comparing their responses to labile-carbon addition in the subsequent bottle-incubation experiment.

### 3.4 Methanogenic potential in selected sediments

Anaerobic bottle incubations showed contrasting patterns in collected gas amounts, methane composition, and soluble chemical oxygen demand (sCOD) between EA3 and EN sediments (Fig. 6). All comparisons are descriptive because each sediment–treatment combination was represented by one bottle. The complete incubation dataset is provided in Table S5.

**Figure 6.**
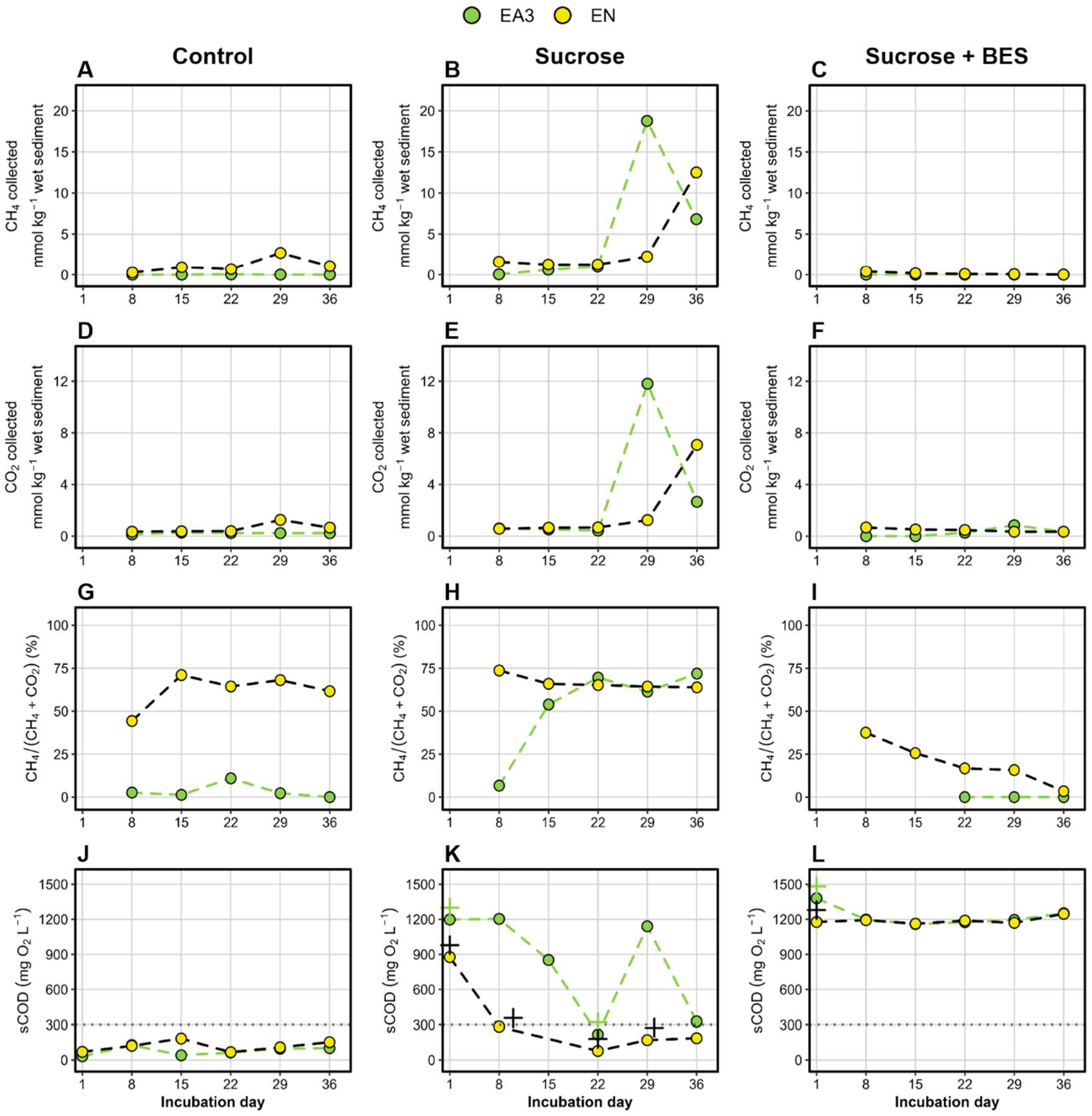
Gas amounts, methane composition, and soluble chemical oxygen demand during sediment incubations. CH_4_ (A–C) and CO_2_ (D–F) amounts, CH_4_ as a percentage of combined measured CH_4_ and CO_2_ (G–I), and soluble chemical oxygen demand (sCOD; J–L) in EA3 (green) and EN (yellow) bottles under unamended control, sucrose, and sucrose + 2-bromoethanesulfonate (BES) treatments. Gas amounts are normalized to initial wet sediment mass and represent successive seven-day collection intervals, plotted at their endpoints on Days 8, 15, 22, 29, and 36. Plus signs indicate sucrose additions; horizontal dotted lines indicate the sCOD feeding threshold of 300 mg O₂ L⁻¹. Normalized CH_4_ is undefined when both measured gases equal zero; these observations are omitted in panel I.

In the controls, collected CH_4_ and CO_2_ amounts were consistently higher in EN than in EA3 (Fig. 6A and D). EN reached maximum interval amounts of 2.7 mmol CH_4_ kg⁻¹ and 1.3 mmol CO_2_ kg⁻¹ wet sediment during Days 22–29, whereas EA3 remained below 0.03 and 0.3 mmol kg⁻¹, respectively. Despite these relatively small gas amounts, CH_4_ accounted for 45–72% of combined measured CH_4_ and CO_2_ in EN, compared with 0–11% in EA3 (Fig. 6G). With sucrose amendment, collected CH_4_ amounts remained below 2 mmol kg⁻¹ in both sediments through Day 22, but gas composition differed early in the incubation (Fig. 6B and H). On Day 8, CH_4_ represented 74% of combined CH_4_ and CO_2_ in EN, compared with 7% in EA3. The EA3 proportion subsequently increased to 54% on Day 15 and approximately 70% on Day 22. The largest collected gas amounts occurred during Days 22–29 in EA3, reaching 18.8 mmol CH_4_ kg⁻¹ and 11.8 mmol CO_2_ kg⁻¹, before declining to 6.8 and 2.7 mmol kg⁻¹ during the final interval. In EN, the highest observed amounts occurred during Days 29–36, reaching 12.5 mmol CH_4_ kg⁻¹ and 7.1 mmol CO_2_ kg⁻¹ (Fig. 6B and E). The sucrose-amended bottles also differed in their sCOD trajectories (Fig. 6K). In EA3, sCOD decreased from approximately 1,200 to 218 mg O₂ L⁻¹ by Day 22, then increased to 1,140 mg O₂ L⁻¹ at the first sampling after sucrose replenishment, on Day 29. In EN, sCOD decreased from 877 to 284 mg O₂ L⁻¹ by Day 8 and remained below 300 mg O₂ L⁻¹ at subsequent measured time points despite repeated sucrose additions. Thus, the sampled EN trajectory showed no comparable sCOD rebound.

In the sucrose + BES bottles, no CH_4_ was detected in EA3, whereas collected CH_4_ in EN declined from 0.40 to 0.01 mmol kg⁻¹ across the incubation (Fig. 6C). The CH_4_ proportion in EN decreased from approximately 38% to 3% of combined CH_4_ and CO_2_ (Fig. 6I). Collected CO_2_ remained below 0.9 mmol kg⁻¹ in both sediments, while sCOD remained high, at approximately 1,160–1,380 mg O₂ L⁻¹ (Fig. 6F and L, respectively).

### 3.6 Community restructuring under sucrose addition and methanogenesis inhibition

Microbial community responses during the bottle incubations were characterized using 16S rRNA gene amplicon sequencing. Family-level composition differed between EA3 and EN and varied across treatments and sampling times (Fig. 7). Control communities shared relatively abundant *Clostridiaceae*, *Anaerolineaceae*, *Bacillaceae*, and *Rhodocyclaceae*, but differed in other bacterial groups. *Geobacteraceae*, *Pseudomonadaceae*, and *Micromonosporaceae* were more prominent in EA3 controls, whereas *Syntrophomonadaceae*, *Syntrophorhabdaceae*, and unresolved assignments within *Desulfotomaculales* were more represented in EN controls.

**Figure 7.**
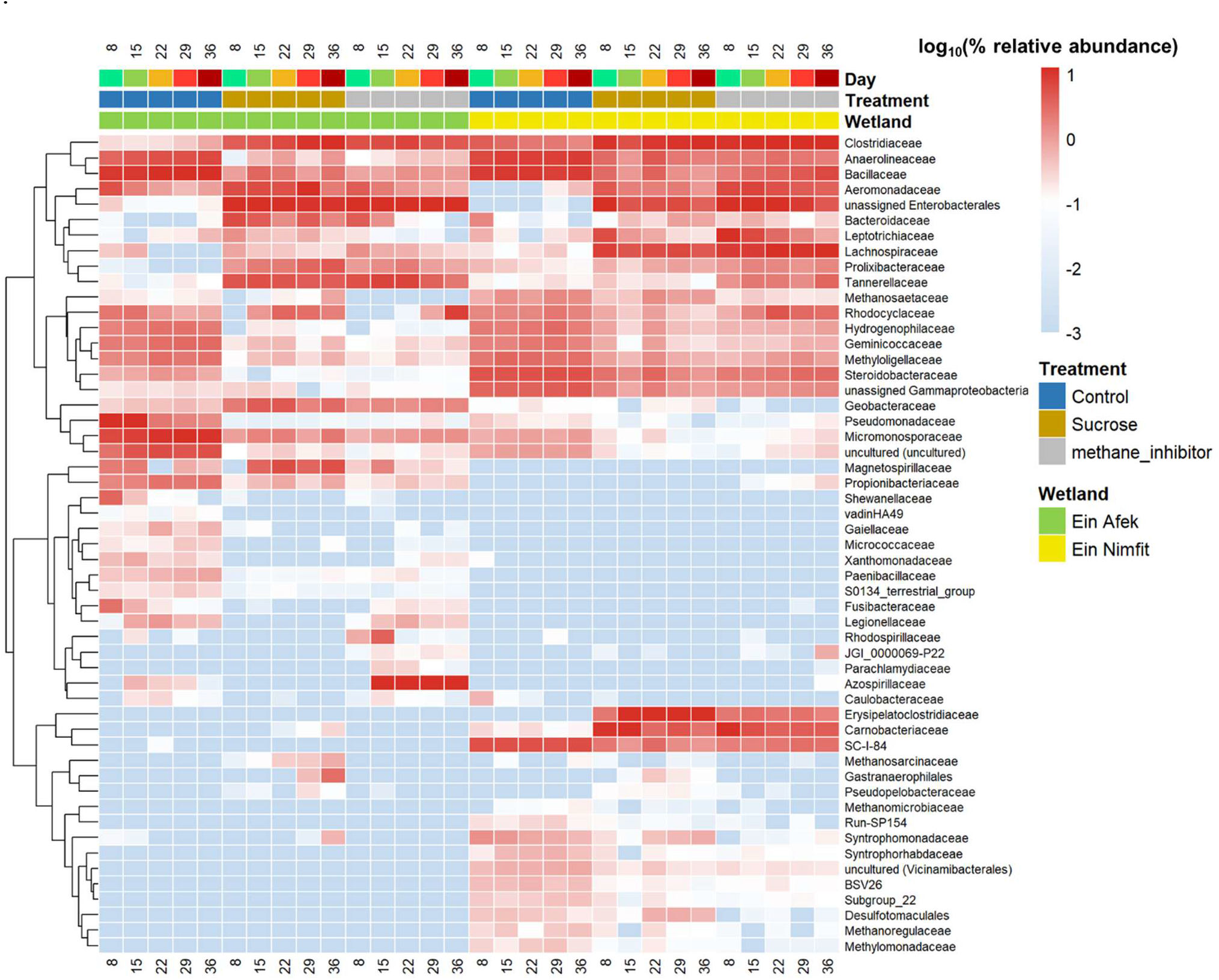
Taxonomic composition of microbial communities during anaerobic sediment incubations. The heatmap shows family-level relative abundances calculated from CSS-normalized 16S rRNA gene counts in EA3 and EN bottles under control, sucrose, and sucrose + BES treatments. Columns represent mean profiles across three technical DNA-extraction replicates for each sediment-treatment-day combination at Days 8, 15, 22, 29, and 36. Rows include the most abundant families and taxa showing the strongest positive or negative Spearman correlations with normalized CH₄ production. Colors represent log₁₀(relative abundance [%] + 0.001). Rows are hierarchically clustered by abundance patterns; columns are ordered by sediment origin, treatment, and sampling day. BES, sodium 2-bromoethanesulfonate.

Sucrose addition was accompanied by increased representation of several shared families, including *Bacteroidaceae*, *Lachnospiraceae*, *Tannerellaceae*, and *Aeromonadaceae*. However, the amended communities retained differences between sediment origins. *Leptotrichiaceae*, *Prolixibacteraceae*, and unresolved family-level assignments within Enterobacterales were more prominent in EA3, whereas *Erysipelatoclostridiaceae* and *Carnobacteriaceae* were more prominent in EN. *Geobacteraceae* showed a particularly contrasting temporal pattern, increasing at later sampling points in the EA3 sucrose bottle while remaining comparatively low in EN. This late increase was less apparent in the EA3 sucrose + BES bottle. The sucrose + BES communities retained several families prominent in the sucrose-only bottles, including *Bacteroidaceae*, *Lachnospiraceae*, *Tannerellaceae*, and *Aeromonadaceae*. Differences between sediments also persisted, with greater representation of *Leptotrichiaceae* in EA3 and *Erysipelatoclostridiaceae* and *Carnobacteriaceae* in EN. These bacterial assemblages persisted despite near-zero CH_4_ production, while continued CO_2_ production indicated that microbial carbon mineralization was maintained under BES treatment (Fig. 6).

Methanogen-associated families exhibited additional variation across bottles and sampling times. *Methanosaetaceae* was comparatively abundant in the controls and increased at later sampling points in the EA3 sucrose bottle, whereas its representation was generally lower under sucrose + BES. *Methanoregulaceae* was also more prominent in the controls than in the amended bottles. *Methanosarcinaceae* and *Methanomicrobiaceae* generally occurred at lower relative abundances; in EA3, *Methanosarcinaceae* was more apparent at early sampling points under sucrose + BES before declining. Methanogen-associated sequences remained detectable in BES-treated bottles despite near-zero CH_4_ production, indicating that their presence did not establish active methanogenesis. Families associated with methylotrophy and methane oxidation, including *Methyloligellaceae* and *Methylomonadaceae*, were also detected, although their activity during the anaerobic incubations was not measured.

Control samples generally had higher Shannon diversity than samples from the amended bottles during the START and END phases, whereas differences in Simpson diversity were less pronounced, particularly in EN (Fig. S2A and B). These phases were defined retrospectively from the observed CH₄ patterns, with the maximum CH₄ percentage in EN used to identify the onset of the END phase. Bray-Curtis NMDS showed separation between EA3 and EN communities and between control and amended communities within each sediment (Fig. S2C). Separation of the control from the sucrose and sucrose + BES communities was more pronounced in EA3 than in EN. Sucrose and sucrose + BES communities overlapped in the ordination, with additional variation across sampling days.

## 4. Discussion

### 4.1 Methane accumulation reflects interactions between sediment carbon, biogeochemistry, and methanogenic communities

High dissolved CH_4_ concentrations occurred in sediments with contrasting organic-carbon and geochemical characteristics (Figs. 2–3). EN contained approximately 12% TOC and reached 2.1 mM CH₄, whereas EA2 contained approximately 5% TOC yet maintained CH_4_ concentrations of 0.7–1.7 mM. Thus, sediment organic-carbon content alone did not explain the observed differences in CH_4_ accumulation. Both CH_4_-rich cores showed substantial SO₄²⁻ depletion with depth, consistent with sulfate reduction. However, EN exhibited pronounced H₂S accumulation and low dissolved Fe, consistent with sulfidic conditions (Plass et al., 2020), whereas EA2 showed comparatively high dissolved Fe and enrichment of NH₄⁺, DIC, and PO₄³⁻. These contrasting profiles suggest differences in the anaerobic processes accompanying CH_4_ accumulation. Competition between sulfate reducers and methanogens for H₂ and acetate could influence substrate availability for methanogenesis (Conrad, 2020; Pester et al., 2012). Nevertheless, dissolved CH_4_ concentrations integrate production, consumption, and transport; they cannot establish how carbon was partitioned among competing pathways. The CH_4_-rich sediments also supported distinct *mcrA* assemblages (Fig. 5). *Methanoregula* dominated the classified community in EA2, consistent with hydrogenotrophic potential supported by fermentation-derived H₂/CO₂ or formate (Fu et al., 2026). EN contained a mixed assemblage of *Methanoregula*, *Methanosarcina*, and *Methanothrix*, suggesting potential utilization of a broader range of methanogenic substrates (Kostopoulou et al., 2023). Although taxonomy does not establish pathway-specific activity, these differences highlight the potential importance of microbial interactions supplying substrates to methanogens (Bechtold et al., 2025; Xing et al., 2026).

In contrast, EA3 combined low CH_4_ concentrations with approximately 1% TOC, low HI, high OI, and persistent SO_4_²⁻. These conditions could constrain methanogenic substrate supply and sustain competition from sulfate reducers (Cheng et al., 2026). Methanogens, including *Methanosarcina*, remained detectable, although Shannon diversity was lowest in EA3 among the sampled cores (Fig. S1). Extensive unresolved *mcrA* assignments limit genus-level interpretation (Cholet et al., 2022; Vasileiadis et al., 2026), but the field profiles indicate that methanogen presence alone did not explain CH_4_ accumulation. This raised a mechanistic question: did EA3 have intrinsically low CH_4_-production capacity, or was methanogenic potential present but constrained by substrate availability under field conditions?

### 4.2 Labile carbon reveals latent methanogenic potential

Sucrose enrichment revealed substantial methanogenic potential in EA3 despite its low field CH_4_ accumulation and lower methanogenic community diversity (Fig. 6; Fig. S1). This observation is consistent with evidence that relationships between methanogen diversity and methane production depend on environmental conditions (Godin et al., 2012), and that responses to organic-carbon enrichment involve the broader microbial community (Bertolet et al., 2022). Both sediments initially yielded small collected methane amounts, but EA3 subsequently exhibited a pronounced, transient increase, whereas EN reached its highest observed output during the final interval. These changes occurred against relatively stable, methane-rich gas composition in EN, while in EA3 the CH_4_ proportion increased before the sharp rise in collected amounts (Fig. 6B and H). Thus, methane enrichment and the magnitude of methane output followed distinct temporal patterns.

The sCOD trajectories suggest that these differences involved carbon processing beyond methanogenesis alone. In EA3, the rebound in sCOD following replenishment coincided with the interval of greatest methane collection. In EN, repeated additions did not produce a comparable accumulation at subsequent sampling points. This raises the possibility that soluble carbon turnover in EN preceded its later increase in methane output. However, sCOD depletion cannot identify the fate of the supplied carbon: uptake into biomass, other respiratory pathways, and methane formation could all contribute. Moreover, fermentation can convert sucrose into soluble organic acids that remain measurable as COD. These observations support considering the development of the broader carbon-processing community alongside methanogens. Sucrose must first be fermented to compounds that methanogens can use directly or following further syntrophic transformations (Conrad, 2020; Hattori, 2008; Yang et al., 2017). Differences in the supply and consumption of these intermediates could contribute to the separation between soluble carbon depletion, methane enrichment, and increased gas output (Valentin-Alvarado et al., 2024).

The contrasting responses of EA3 and EN may reflect their initial microbial communities together with differences in sediment carbon pools and geochemical conditions. EA3 contained approximately 1% TOC, with low HI and high OI, whereas EN contained approximately 12% TOC and comparatively high HI. The same amendment was therefore introduced into substantially different sediment environments. Previous incubation and stable-isotope-probing studies similarly highlight interactions between methanogens and other anaerobic microorganisms in the expression of methanogenic potential (Hamovit et al., 2025). The transient EA3 response could reflect depletion of sucrose or fermentation-derived intermediates; however, neither was measured, preventing identification of the mechanism underlying the subsequent decline in gas production. These responses describe the individual bottles examined, given the unreplicated incubation design.

The strong EA3 response suggests that sediments with low field CH_4_ accumulation may nevertheless respond markedly to fresh carbon inputs. In natural wetlands, root exudates and litter deposition can supply substrates supporting anaerobic carbon processing and methanogenesis (Berrittella & Van Huissteden, 2011), while changes in flooding duration and water residence time can alter oxygen exposure and electron-acceptor availability (Tokarz & Urban, 2015). Together, these drivers may influence the expression of methanogenic potential under changing wetland conditions.

### 4.3 Carbon enrichment reveals distinct routes to methane production potential

Sucrose amendment was accompanied by changes in alpha diversity and community composition that differed between EA3 and EN (Fig. S2). EA3 showed more pronounced community restructuring alongside its transient CH_4_ response, whereas EN showed a more gradual gas response and different compositional changes. Similar observations in wetland microbiomes show that taxonomically distinct communities can respond differently to environmental perturbations while retaining overlapping carbon-processing functions (Zhang et al., 2025). The unamended bottles show that EA3 and EN differed in the anaerobic communities present before any sucrose addition (Fig. 7). EN contained higher relative abundances of *Methanosaetaceae* and *Methanoregulaceae*, together with *Syntrophomonadaceae* and *Syntrophorhabdaceae*. The latter two families include organisms that can degrade organic intermediates in partnership with methanogens (McInerney et al., 2009; Qiu et al., 2008). Their co-occurrence in the EN control is consistent with a community capable of sustaining methanogenesis on substrates already present in the sediment, which could help explain why the collected gas was more methane rich in EN even without sucrose. In contrast, the EA3 control showed greater representation of *Pseudomonadaceae* and *Micromonosporaceae* and less prominent representation of these putative methanogenic partnerships. These profiles identify differences in community composition that could contribute to the control gas contrast, although they do not establish which organisms supplied substrates or produced methane.

Sucrose amendment was accompanied by greater compositional restructuring in EA3 than in EN (Figs. 7 and S2). Across the amended bottles, several bacterial groups with members previously associated with fermentation, facultative anaerobic metabolism, or organic carbon degradation became more prominent. These included *Bacteroidaceae*, *Lachnospiraceae*, *Tannerellaceae*, *Prolixibacteraceae*, *Aeromonadaceae*, and unassigned *Enterobacterales* (Hamberger et al., 2008; Maus et al., 2022; Steiniger et al., 2023; Zhu et al., 2024). In EA3, *Bacteroidaceae*, *Tannerellaceae*, and unassigned *Enterobacterales* were already more prominent in the sucrose bottle than in the control by Day 8, whereas *Methanosaetaceae* and *Methanosarcinaceae* became more prominent at later sampling points. The methane fraction also rose before the peak in collected CH₄. Together, these patterns are consistent with bacterial processing of added carbon followed by changes in the supply and use of fermentation products by methanogens (McInerney et al., 2009; Stams & Plugge, 2009). However, family-level 16S rRNA gene profiles cannot establish which taxa were active or which substrates they used.

In EN, the amended community was distinguished by prominent Lachnospiraceae, Leptotrichiaceae, Erysipelatoclostridiaceae, and Carnobacteriaceae, alongside bacterial groups shared with amended EA3. These contrasting profiles suggest that the two sediments differed in how their microbial communities responded to fresh carbon, although family-level relative abundances cannot establish which taxa supplied or consumed methanogenic substrates. The EN bottle maintained a relatively stable, methane-rich gas composition even as the amount collected changed. Together, the two trajectories suggest that fresh carbon can support methane formation through different community responses: a pronounced shift in the representation of potential substrate suppliers and methanogens in EA3, and changing methane output without a comparable shift in gas composition in EN. Yet, family-level assignments and relative abundances cannot establish the pathways or rates underlying either response.

The sucrose + BES bottles provide a useful check on the above observations. Several bacterial groups prominent under sucrose alone remained prominent with BES, including unassigned *Enterobacterales* in both sediments, *Bacteroidaceae* and *Tannerellaceae* in EA3, and *Lachnospiraceae* and *Leptotrichiaceae* in EN. Yet collected CH_4_ remained low. Thus, the bacterial response to sucrose was observable even when methane output was small. The BES treatment also cannot be interpreted as a selective subtraction of methane from an otherwise unchanged system: reduced methanogenic consumption of hydrogen can alter the conditions for upstream metabolism and change bacterial community composition, as shown in anaerobic bioreactors (Shrestha et al., 2023). The differences between the sucrose and sucrose + BES profiles are therefore compatible with feedback between methanogens and other anaerobes, but hydrogen and fermentation intermediates were not measured here.

The clearest implication is that low field CH_4_ accumulation and lower methanogen diversity did not preclude substantial methane production in EA3 after sucrose enrichment. The field profiles identified sediment conditions associated with CH_4_ accumulation, while the incubations revealed contrasting responses to fresh carbon in EA3 and EN. In both sediments, bacterial community shifts were evident before the largest amounts of CH_4_ were collected. This timing suggests a testable hypothesis: the methane response to fresh carbon depends on how closely the production of fermentation intermediates is coupled to their consumption by methanogens. Tracking these intermediates alongside microbial activity would help determine what governs the timing of methane production after carbon enrichment.

## 5. Conclusions

Methane accumulation across the sampled wetland sediments was associated with distinct combinations of organic-carbon properties, porewater chemistry, and methanogenic community composition. High CH_4_ concentrations occurred in both organic-carbon-rich EN and moderately carbon-rich EA2, despite contrasting sulfide, iron, nutrient, and classified methanogen profiles. Conversely, increasing organic-carbon content with depth in EA1 did not coincide with substantial CH_4_ accumulation. These findings show that sediment carbon quantity alone was insufficient to account for the observed methane distribution and underscore the importance of its biogeochemical context. Complementary incubations revealed substantial methanogenic potential in EA3 despite its low field CH_4_ accumulation, low sediment organic-carbon content, and lower methanogenic diversity. Contrasting gas composition, collected amounts, sCOD patterns, and community changes in EA3 and EN suggest differences in the development of anaerobic carbon processing. Although the unreplicated experiment cannot resolve the mechanisms responsible, it demonstrates the value of examining functional responses alongside field profiles. Together, these approaches distinguish methane accumulation, microbial presence, and expressed production potential, supporting an integrated assessment of carbon properties, sediment chemistry, and broader microbial communities in wetland methane research.

## References

Alewell, C., Paul, S., Lischeid, G., & Storck, F. R. (2008). Co-regulation of redox processes in freshwater wetlands as a function of organic matter availability? Science of the Total Environment, 404(2–3), 335–342.

Bechtold, E. K., Ellenbogen, J. B., Villa, J. A., De Melo Ferreira, D. K., Oliverio, A. M., Kostka, J. E., Rich, V. I., Varner, R. K., Bansal, S., Ward, E. J., Bohrer, G., Borton, M. A., Wrighton, K. C., & Wilkins, M. J. (2025). Metabolic interactions underpinning high methane fluxes across terrestrial freshwater wetlands. Nature Communications, 16(1), 944. 10.1038/s41467-025-56133-0

Berberich, M. E., Beaulieu, J. J., Hamilton, T. L., Waldo, S., & Buffam, I. (2020). Spatial variability of sediment methane production and methanogen communities within a eutrophic reservoir: Importance of organic matter source and quantity. Limnology and Oceanography, 65(6), 1336–1358. 10.1002/lno.11392

Berrittella, C., & Van Huissteden, J. (2011). Uncertainties in modelling CH_4_ emissions from northern wetlands in glacial climates: The role of vegetation parameters. Climate of the Past, 7(4), 1075–1087. 10.5194/cp-7-1075-2011

Bertolet, B. L., West, W. E., Armitage, D. W., & Jones, S. E. (2019). Organic matter supply and bacterial community composition predict methanogenesis rates in temperate lake sediments. Limnology and Oceanography Letters, 4(5), 164–172. 10.1002/lol2.10114

Bertolet, B. L., Koepfli, C., & Jones, S. E. (2022). Lake sediment methane responses to organic matter are related to microbial community composition in experimental microcosms. Frontiers in Environmental Science, 10, 834829. 10.3389/fenvs.2022.834829

Biancalani, R., Avagyan, A., & Food and Agriculture Organization of the United Nations (Eds.). (2014). Towards climate-responsible peatlands management. Food and Agriculture Organization of the United Nations (FAO).

Bolyen, E., Rideout, J. R., Dillon, M. R., Bokulich, N. A., Abnet, C. C., Al-Ghalith, G. A., Alexander, H., Alm, E. J., Arumugam, M., Asnicar, F., Bai, Y., Bisanz, J. E., Bittinger, K., Brejnrod, A., Brislawn, C. J., Brown, C. T., Callahan, B. J., Caraballo-Rodríguez, A. M., Chase, J., … Caporaso, J. G. (2019). Reproducible, interactive, scalable and extensible microbiome data science using QIIME 2. Nature Biotechnology, 37(8), 852–857. 10.1038/s41587-019-0209-9

Bueno De Mesquita, C. P., Wu, D., & Tringe, S. G. (2023). Methyl-based methanogenesis: An ecological and genomic review. Microbiology and Molecular Biology Reviews, 87(1), e00024–22. 10.1128/mmbr.00024-22

Callahan, B. J., McMurdie, P. J., Rosen, M. J., Han, A. W., Johnson, A. J. A., & Holmes, S. P. (2016). DADA2: High-resolution sample inference from Illumina amplicon data. Nature Methods, 13(7), 581–583. 10.1038/nmeth.3869

Cheng, T., Pan, W., Wanek, W., Tan, C., Bol, R., Liang, Y., & Ma, Q. (2026). From substrate limitation to competition control: Straw input regulates sulfate-mediated suppression of methanogenesis in paddy soils. Soil Biology and Biochemistry, 222, 110263. 10.1016/j.soilbio.2026.110263

Cholet, F., Lisik, A., Agogué, H., Ijaz, U. Z., Pineau, P., Lachaussée, N., & Smith, C. J. (2022). Ecological observations based on functional gene sequencing are sensitive to the amplicon processing method. Msphere, 7(4), e00324–22.

Cline, J. D. (1969). Spectrophotometric determination of hydrogen sulfide in natural waters 1. Limnology and Oceanography, 14(3), 454–458. 10.4319/lo.1969.14.3.0454

Conrad, R. (2020). Importance of hydrogenotrophic, aceticlastic and methylotrophic methanogenesis for methane production in terrestrial, aquatic and other anoxic environments: A mini review. Pedosphere, 30(1), 25–39. 10.1016/S1002-0160(18)60052-9

Dalcin Martins, P., Hoyt, D. W., Bansal, S., Mills, C. T., Tfaily, M., Tangen, B. A., Finocchiaro, R. G., Johnston, M. D., McAdams, B. C., Solensky, M. J., Smith, G. J., Chin, Y., & Wilkins, M. J. (2017). Abundant carbon substrates drive extremely high sulfate reduction rates and methane fluxes in Prairie Pothole Wetlands. Global Change Biology, 23(8), 3107–3120. 10.1111/gcb.13633

Deshmukh, C. S., Julius, D., Desai, A. R., Asyhari, A., Page, S. E., Nardi, N., Susanto, A. P., Nurholis, N., Hendrizal, M., Kurnianto, S., Suardiwerianto, Y., Salam, Y. W., Agus, F., Astiani, D., Sabiham, S., Gauci, V., & Evans, C. D. (2021). Conservation slows down emission increase from a tropical peatland in Indonesia. Nature Geoscience, 14(7), 484–490. 10.1038/s41561-021-00785-2

Disnar, J. R., Guillet, B., Keravis, D., Di-Giovanni, C., & Sebag, D. (2003). Soil organic matter (SOM) characterization by Rock-Eval pyrolysis: Scope and limitations. Organic Geochemistry, 34(3), 327–343. 10.1016/S0146-6380(02)00239-5

Ferry, J. G. (2010). The chemical biology of methanogenesis. Planetary and Space Science, 58(14–15), 1775–1783. 10.1016/j.pss.2010.08.014

Ferry, J. G. (2011). Fundamentals of methanogenic pathways that are key to the biomethanation of complex biomass. Current Opinion in Biotechnology, 22(3), 351–357. 10.1016/j.copbio.2011.04.011

Frolking, S., Talbot, J., Jones, M. C., Treat, C. C., Kauffman, J. B., Tuittila, E.-S., & Roulet, N. (2011). Peatlands in the Earth’s 21st century climate system. Environmental Reviews, 19(NA), 371–396. 10.1139/a11-014

Fu, M., Liu, Y., Li, P., Yi, X., Liu, Y., Zhan, J., & Zhou, H. (2026). Metagenomic mechanisms of acidification control by FeOx@HCS in stabilizing methanogenesis. Biochemical Engineering Journal, 229, 110110. 10.1016/j.bej.2026.110110

Gerera, Y., Pellerin, A., Eliani Russak, E., Walter Anthony, K., Hasson, N., Oved Rosenberg, Y., & Sivan, O. (2025). The evolution of methane production rates from young to mature thermokarst lakes. Biogeosciences, 22(23), 7901–7914. 10.5194/bg-22-7901-2025

Godin, A., McLaughlin, J. W., Webster, K. L., Packalen, M., & Basiliko, N. (2012). Methane and methanogen community dynamics across a boreal peatland nutrient gradient. Soil Biology and Biochemistry, 48, 96–105. 10.1016/j.soilbio.2012.01.018

Goldstein, A., Turner, W. R., Spawn, S. A., Anderson-Teixeira, K. J., Cook-Patton, S., Fargione, J., Gibbs, H. K., Griscom, B., Hewson, J. H., Howard, J. F., Ledezma, J. C., Page, S., Koh, L. P., Rockström, J., Sanderman, J., & Hole, D. G. (2020). Protecting irrecoverable carbon in Earth’s ecosystems. Nature Climate Change, 10(4), 287–295. 10.1038/s41558-020-0738-8

Hamberger, A., Horn, M. A., Dumont, M. G., Murrell, J. C., & Drake, H. L. (2008). Anaerobic consumers of monosaccharides in a moderately acidic fen. Applied and Environmental Microbiology, 74(10), 3112–3120. 10.1128/AEM.00193-08

Hamovit, N., RoyChowdhury, T., Akob, D. M., Zhang, X., McCarty, G., & Yarwood, S. (2025). Comparative assessment of a restored and natural wetland using^13^ C-DNA SIP reveals a higher potential for methane production in the restored wetland. Applied and Environmental Microbiology, 91(3), e02161–24. 10.1128/aem.02161-24

Hartman, W. H., Bueno De Mesquita, C. P., Theroux, S. M., Morgan-Lang, C., Baldocchi, D. D., & Tringe, S. G. (2024). Multiple microbial guilds mediate soil methane cycling along a wetland salinity gradient. mSystems, 9(1), e00936–23. 10.1128/msystems.00936-23

Hattori, S. (2008). Syntrophic acetate-oxidizing microbes in methanogenic environments. Microbes and Environments, 23(2), 118–127. 10.1264/jsme2.23.118

Hazera, J., Sebag, D., Kowalewski, I., Ravelojaona, H., Verrecchia, E., Jakab, G., Zacháry, D., Schneider, F., Trombino, L., Manlay, R. J., Fouché, J., & Chevallier, T. (2026). Technical note: Further adjustments to the Rock-Eval® thermal analysis for soil organic and inorganic carbon quantification to avoid post-hoc corrections. Biogeosciences, 23(5), 1881–1895. 10.5194/bg-23-1881-2026

Kayranli, B., Scholz, M., Mustafa, A., & Hedmark, Å. (2010). Carbon storage and fluxes within freshwater wetlands: A critical review. Wetlands, 30, 111–124.

Kostopoulou, E., Chioti, A. G., Tsioni, V., & Sfetsas, T. (2023). Microbial dynamics in anaerobic digestion: A review of operational and environmental factors affecting microbiome composition and function. 10.20944/preprints202306.0299.v1

Laanbroek, H. J. (2010). Methane emission from natural wetlands: Interplay between emergent macrophytes and soil microbial processes. A mini-review. Annals of Botany, 105(1), 141–153.

Lehmann, J., & Kleber, M. (2015). The contentious nature of soil organic matter. Nature, 528(7580), 60–68. 10.1038/nature16069

Liu, J., Klonicki-Ference, E., Krause, S. J. E., & Treude, T. (2025). Iron oxides fuel anaerobic oxidation of methane inthe presence of sulfate in hypersaline coastal wetland sediment. Environmental Science & Technology, 59(1), 513–522. 10.1021/acs.est.4c10639

Lorenz, K., & Lal, R. (2018). Carbon sequestration in wetland soils. In K. Lorenz & R. Lal, Carbon Sequestration in Agricultural Ecosystems (pp. 211–234). Springer International Publishing. 10.1007/978-3-319-92318-5_5

Luton, P. E., Wayne, J. M., Sharp, R. J., & Riley, P. W. (2002). The mcrA gene as an alternative to 16S rRNA in the phylogenetic analysis of methanogen populations in landfill. Microbiology, 148(11), 3521–3530. 10.1099/00221287-148-11-3521

Marschner, B., & Kalbitz, K. (2003). Controls of bioavailability and biodegradability of dissolved organic matter in soils. Geoderma, 113(3–4), 211–235. 10.1016/S0016-7061(02)00362-2

Martin, M. (2011). Cutadapt removes adapter sequences from high-throughput sequencing reads. EMBnet.journal, 17(1), 10–12. 10.14806/ej.17.1.200

Maus, I., Wibberg, D., Belmann, P., Hahnke, S., Huang, L., Spröer, C., Bunk, B., Blom, J., Sczyrba, A., Pühler, A., Klocke, M., & Schlüter, A. (2022). The novel oligopeptide utilizing species Anaeropeptidivorans aminofermentans M3/9T, its role in anaerobic digestion and occurrence as deduced from large-scale fragment recruitment analyses. Frontiers in Microbiology, 13, 1032515. 10.3389/fmicb.2022.1032515

McInerney, M. J., Sieber, J. R., & Gunsalus, R. P. (2009). Syntrophy in anaerobic global carbon cycles. Current Opinion in Biotechnology, 20(6), 623–632. 10.1016/j.copbio.2009.10.001

Megonigal, J. P., Hines, M. E., & Visscher, P. T. (2003). Anaerobic metabolism: Linkages to trace gases and aerobic processes. In Treatise on Geochemistry (pp. 317–424). Elsevier. 10.1016/B0-08-043751-6/08132-9

Naqib, A., Poggi, S., Wang, W., Hyde, M., Kunstman, K., & Green, S. J. (2018). Making and sequencing heavily multiplexed, high-throughput 16S ribosomal RNA gene amplicon libraries using a flexible, two-stage PCR protocol. In N. Raghavachari & N. Garcia-Reyero (Eds.), Gene Expression Analysis (Vol. 1783, pp. 149–169). Springer New York. 10.1007/978-1-4939-7834-2_7

Pester, M., Knorr, K.-H., Friedrich, M. W., Wagner, M., & Loy, A. (2012). Sulfate-reducing microorganisms in wetlands–fameless actors in carbon cycling and climate change. Frontiers in Microbiology, 3, 72.

Plass, A., Schlosser, C., Sommer, S., Dale, A. W., Achterberg, E. P., & Scholz, F. (2020). The control of hydrogen sulfide on benthic iron and cadmium fluxes in the oxygen minimum zone off Peru. Biogeosciences, 17(13), 3685–3704. 10.5194/bg-17-3685-2020

Qiu, Y.-L., Hanada, S., Ohashi, A., Harada, H., Kamagata, Y., & Sekiguchi, Y. (2008). Syntrophorhabdus aromaticivorans gen. nov., sp. nov., the first cultured anaerobe capable of degrading phenol to acetate in obligate syntrophic associations with a hydrogenotrophic methanogen. Applied and Environmental Microbiology, 74(7), 2051–2058. 10.1128/AEM.02378-07

Reddy, K. R., DeLaune, R. D., & Inglett, P. W. (2022). Biogeochemistry of wetlands: Science and applications. CRC press.

Reiche, M., Torburg, G., & KÃ¼sel, K. (2008). Competition of Fe(III) reduction and methanogenesis in an acidic fen: Fe(III) reduction and methanogenesis in a fen. FEMS Microbiology Ecology, 65(1), 88–101. 10.1111/j.1574-6941.2008.00523.x

Sapir, G., & Elron, E. (2014). Water plan for the Na‘aman River: Final report. Israel Nature and Parks Authority and Western Galilee Drainage Authority. (In Hebrew).

Sapir, G., Angert, A., Rosenberg, Y. O., & Golan, R. (2026). Long-term effects of drainage and rewetting on the degradation and preservation of peat organic matter in sub-tropical climate. Biogeosciences, 23(4), 1403–1421. 10.5194/bg-23-1403-2026

Schmidt, M. W. I., Torn, M. S., Abiven, S., Dittmar, T., Guggenberger, G., Janssens, I. A., Kleber, M., Kögel-Knabner, I., Lehmann, J., Manning, D. A. C., Nannipieri, P., Rasse, D. P., Weiner, S., & Trumbore, S. E. (2011). Persistence of soil organic matter as an ecosystem property. Nature, 478(7367), 49–56. 10.1038/nature10386

Scott, B., Baldwin, A. H., Ballantine, K., Palmer, M., & Yarwood, S. (2020). The role of organic amendments in wetland restorations. Restoration Ecology, 28(4), 776–784. 10.1111/rec.13179

Sebag, D., Verrecchia, E. P., Cécillon, L., Adatte, T., Albrecht, R., Aubert, M., Bureau, F., Cailleau, G., Copard, Y., Decaens, T., Disnar, J.-R., Hetényi, M., Nyilas, T., & Trombino, L. (2016). Dynamics of soil organic matter based on new Rock-Eval indices. Geoderma, 284, 185–203. 10.1016/j.geoderma.2016.08.025

Sela-Adler, M., Ronen, Z., Herut, B., Antler, G., Vigderovich, H., Eckert, W., & Sivan, O. (2017). Co-existence of methanogenesis and sulfate reduction with common substrates in sulfate-rich estuarine sediments. Frontiers in Microbiology, 8, 766. 10.3389/fmicb.2017.00766

Shrestha, S., Xue, S., & Raskin, L. (2023). Competitive reactions during ethanol chain elongation were temporarily suppressed by increasing hydrogen partial pressure through methanogenesis inhibition. Environmental Science & Technology, 57(8), 3369–3379. 10.1021/acs.est.2c09014

Stams, A. J. M., & Plugge, C. M. (2009). Electron transfer in syntrophic communities of anaerobic bacteria and archaea. Nature Reviews Microbiology, 7(8), 568–577. 10.1038/nrmicro2166

Steiniger, B., Hupfauf, S., Insam, H., & Schaum, C. (2023). Exploring anaerobic digestion from mesophilic to thermophilic temperatures—Operational and microbial aspects. Fermentation, 9(9), 798. 10.3390/fermentation9090798

Stookey, L. L. (1970). Ferrozine—A new spectrophotometric reagent for iron. Analytical Chemistry, 42(7), 779–781. 10.1021/ac60289a016

Takahashi, S., Tomita, J., Nishioka, K., Hisada, T., & Nishijima, M. (2014). Development of a prokaryotic universal primer for simultaneous analysis of bacteria and archaea using next-generation sequencing. PLoS ONE, 9(8), e105592. 10.1371/journal.pone.0105592

Tapio-Biström, M.-L., Joosten, H., Tol, S., Mitigation of Climate Change in Agriculture Programme, & Wetlands International (Eds.). (2012). Peatlands: Guidance for climate change mitigation through conservation, rehabilitation and sustainable use (2nd ed). Food and Agriculture Organization of the United Nations: Wetlands International.

Tokarz, E., & Urban, D. (2015). Soil redox potential and its impact on microorganisms and plants of wetlands. Journal of Ecological Engineering, 16, 20–30. 10.12911/22998993/2801

Valentin-Alvarado, L. E., Appler, K. E., De Anda, V., Schoelmerich, M. C., West-Roberts, J., Kivenson, V., Crits-Christoph, A., Ly, L., Sachdeva, R., Greening, C., Savage, D. F., Baker, B. J., & Banfield, J. F. (2024). Asgard archaea modulate potential methanogenesis substrates in wetland soil. Nature Communications, 15(1), 6384. 10.1038/s41467-024-49872-z

Vasileiadis, S., Valmas, M. I., Pitsikoglou, D. S., Rodosthenous, J., Omirou, M., Kotsopoulos, T. A., Yan, Y., Fu, D., & Fotidis, I. A. (2026). Up-to-date, and taxonomy-curated mcrA reference databases for methanogen community profiling. Systematic and Applied Microbiology, 126752.

Waldo, N. B., Hunt, B. K., Fadely, E. C., Moran, J. J., & Neumann, R. B. (2019). Plant root exudates increase methane emissions through direct and indirect pathways. Biogeochemistry, 145(1–2), 213–234. 10.1007/s10533-019-00600-6

Xing, T., Ji, M., Liu, K., Liu, P., & Liu, Y. (2026). Elevated microbial network complexity and enriched methanogens contribute to higher methanogenic potential in hollows than in hummocks in alpine meadows. Environmental Microbiome, 21(1), 40. 10.1186/s40793-026-00859-x

Yang, B., Wang, Q., Zhao, X., Wang, C., Liu, J., Yang, H., Liu, S., Yin, F., & Zhang, W. (2017). Increased energy production from sucrose by controlled hydrogen-producing fermentation followed by methanogenic fermentation. Energy Sources, Part A: Recovery, Utilization, and Environmental Effects, 39(6), 576–582. 10.1080/15567036.2016.1246627

Yarwood, S. A. (2018). The role of wetland microorganisms in plant-litter decomposition and soil organic matter formation: A critical review. FEMS Microbiology Ecology, 94(11). 10.1093/femsec/fiy175

Yusim, E. J., Zarecki, R., Medina, S., Carmi, G., Mousa, S., Hassanin, M., Ronen, Z., Wu, Z., Jiang, J., Baransi-Karkaby, K., Avisar, D., Sabbah, I., Yanuka-Golub, K., & Freilich, S. (2025). Integrated use of electrochemical anaerobic reactors and genomic based modeling for characterizing methanogenic activity in microbial communities exposed to BTEX contamination. Environmental Research, 268, 120691. 10.1016/j.envres.2024.120691

Zhang J., Kobert K., Flouri T., Stamatakis A. (2014). PEAR: a fast and accurate Illumina Paired-End reAd mergeR. Bioinformatics 30:614–620. 10.1093/bioinformatics/btt593.

Zhang, H., Zhao, D., Wu, Q. L., & Zeng, J. (2025). Environment selected microbial function rather than taxonomic species in a plateau saline-alkaline wetland. Applied and Environmental Microbiology, 91(7), e02206–24. 10.1128/aem.02206-24

Zheng, H., Liu, D., Li, Y., Chen, Z., Li, J., Dong, Y., Yang, C., Miao, Y., Yuan, J., & Ding, W. (2025). Wetland restoration suppresses microbial carbon metabolism by altering keystone species interactions. Frontiers in Microbiology, 16, 1570703. 10.3389/fmicb.2025.1570703

Zhu, X., Li, P., & Ju, F. (2024). Microbiome dynamics and products profiles of biowaste fermentation under different organic loads and additives. Engineering in Life Sciences, 24(5), 2300216. 10.1002/elsc.202300216

